# Hemodynamic Analysis of a Repaired Ascending Aorta with Preserved Aortic Root

**DOI:** 10.64898/2026.01.28.702307

**Authors:** Hannah Zhai, Yurui Chen, Yuichiro Kitada, Hiroo Takayama, Vijay Vedula

## Abstract

**Purpose:** To evaluate the hemodynamic impact of restoring a normal sino-tubular junction (STJ) following a novel Hegar dilator-based procedure in patients undergoing root-sparing ascending thoracic aortic aneurysm (ATAA) repair using computational modeling.

**Methods:** We retrospectively selected an ATAA patient who underwent pre- and postoperative gated computed tomography angiography (CTA). We developed a novel workflow to segment the lumen, thick-walled aorta, and aortic valve from CTA images for subsequent blood flow analysis using computational fluid dynamics (CFD) and fluid-structure interaction (FSI). Morphological and hemodynamic characteristics of the root were quantified and compared against those of a control subject, with no noted ascending aortic dilation. The model’s sensitivity to graft properties and leaflet material heterogeneity was analyzed.

**Results:** Both CFD and FSI results showed that the postoperative geometry reconstructed with a normal STJ profile reintroduces sinus vortices during peak systole, similar to the control subject, but were absent pre-surgery. Accounting for aortic valve leaflets in FSI studies yielded qualitatively similar results to the CFD cases, albeit with locally elevated velocities, time-averaged wall shear stress (TAWSS), and energy dissipation, likely due to the dynamically changing orifice area and differing profiles of the left ventricular outflow tract (LVOT).

**Conclusion:** We demonstrated that the novel Hegar dilator-based STJ reconstruction restores normal blood flow patterns, highlighting the importance of reprofiling the aortic sinuses and STJ. The study also highlights the model’s sensitivities, particularly the LVOT shape and leaflet morphology and mobility, and may assist planning STJ reconstruction to yield optimal hemodynamics before intervention.

## 1 Introduction

Thousands of patients with ascending thoracic aortic aneurysms (ATAA) undergo surgical replacement every year to mitigate the risk of life-threatening events, including aortic dissection or rupture [1]. The threshold for surgical intervention is primarily guided by the diameter of the aortic lumen exceeding 5 cm [2, 3], although recent clinical studies advocate creating separate thresholds for the aortic root^1^ and the distal ascending aorta [4]. An area of particular interest is the sino-tubular junction (STJ), located between the aortic root and ascending aorta, which plays an essential role in the normal functioning of the aortic valve [5]. An aneurysmal STJ region or aortic root is at risk of progressive root dilation leading to aortic regurgitation, in which improper aortic valve closure causes the blood to flow back into the left ventricle [5–8].

Several phenotypical variants of ATAA exist, including an enlarged aortic root, STJ, and ascending aorta, which manifest either individually or in combination [9]. While ascending aortic replacement remains the most commonly performed surgical procedure for ATAA [10], other strategies are available to treat ATAA patients depending on the phenotype. For instance, the Bentall procedure is performed when both the ascending aorta and the aortic root are aneurysmal in conjunction with a faulty regurgitating aortic valve. Here, the entire ascending aorta is replaced with a conduit, and a prosthetic valve (bioprosthetic or mechanical) is implanted in place of the faulty native valve [11, 12]. Alternatively, a valve-sparing ascending aortic replacement surgery, known as the David procedure [13], seeks to retain the native valve, thereby avoiding the complications associated with prosthetic valves used in the Bentall procedure, such as calcifications or blood clots necessitating lifelong use of anticoagulants [14]. In cases with a dilated STJ but normal root, a root surgery is not recommended as replacing the whole ascending aorta with the root carries unnecessary operative risks [15]. For these patients, STJ dilation is treated using an ascending aortic replacement procedure that maintains the native aortic root, including the native valve. Therefore, only the aneurysmal portion of the ascending aorta is replaced, and the naturally occurring STJ constriction, as seen in healthy subjects, is restored. Despite these procedural advances in treating segmentally or wholly dilated ascending aorta, a standardized approach for STJ reconstruction has not been established.

Our team of surgeons at Columbia University Medical Center (CUMC) has developed a novel technique to treat patients with ATAA who have dilated STJ and an intact aortic valve. We note that the details of the procedure and its clinical merits will be published in a peer-reviewed surgical journal and are not the focus of this study. Here, we will only provide a brief overview of this technique to give the reader sufficient background and motivate the current work. The overall goal of the procedure is to restore a biomimetic aortic root and STJ morphology by resizing the STJ and reprofiling the shape and size of the aortic sinuses, bespoke to the patient, while mimicking a normal aortic cusp’s profile. This procedure utilizes a Hegar dilator to tighten a prosthetic Dacron graft in the STJ region to a patient-specific optimal size determined from preoperative computerized tomography angiography (CTA) imaging. After tightening, the graft is sutured to the native aorta.

Compared to previously reported strategies for treating dilated STJ [16–18], this method benefits from the selection and implementation of a subject-specific diameter for the reconstructed STJ, taking into consideration the patient’s individual factors such as the diameter of the basal ring, Sinus of Valsalva, and the operating surgeon’s experience. Postoperative imaging confirmed that the reconstructed geometry is more representative of a healthy aorta, characterized by an apparent constriction at the STJ and rounded aortic sinus profiles. However, it remains unclear whether restoring a normal aortic root and STJ profile is adequate to yield normal blood flow patterns and hemodynamic forces in the root region. Examining flow patterns in the aortic root is particularly important in cases of STJ effacement, as an altered hemodynamic and mechanical environment in the aortic root and valve can trigger a root aneurysm, an intimal tear leading to a type A aortic dissection, or valve degeneration. Furthermore, it raises the possibility of using predictive computational modeling to determine the hemodynamically optimal graft shape and size, thereby preplanning surgery tailored to the patient through virtual deployment, which complements the current imaging-only approach [19].

In this study, we aim to evaluate the hemodynamic efficacy of the new STJ reconstruction procedure through a comprehensive computational analysis of blood flow in the ascending aorta of a retrospectively selected patient with ATAA. Specifically, we adopted a two-pronged approach to simulate hemodynamics in the thoracic aorta, constructed from the patient’s gated CTA images, acquired before and after the surgery. First, we assumed the aortic wall to be rigid and performed a computational fluid dynamics (CFD) simulation of blood flow through the thoracic aorta. Second, we considered the aorta to be deformable and incorporated the aortic valve to simulate the interactions between blood, the aortic valve, and the deformable aortic wall using a fluid-structure interaction (FSI) model. Throughout this study, we employed SimVascular’s image segmentation pipeline [20, 21] to construct the patient-specific finite element analysis (FEA) model of the thoracic aorta. We also developed a novel workflow to construct a patient-specific aortic valve from CTA images, fused into the aortic wall as a single block of tissue, but with different material parameters for the valve and the vessel. In both CFD and FSI cases, we compared the aortic root hemodynamics, including blood flow patterns, shear forces, and kinetic energy budget, before and after the surgery in the ATAA patient, and compared these measures against those of a control subject. The FSI model was also used to analyze leaflet opening dynamics, followed by a sensitivity analysis of the vessel and valve leaflet material parameters.

The manuscript is structured as follows: Section 2 provides a detailed description of the study design, including simulation workflow and methods employed. This includes patient data, the image-to-model creation pipeline for both CFD and FSI studies, an overview of governing equations, boundary conditions, and a description of the FEA solver and simulation setup. Section 3 highlights key results obtained from the CFD and FSI studies, followed by a detailed discussion of the results, model sensitivities, study limitations, and conclusions in Section 4.

## 2 Materials and Methods

### 2.1 Patient Information

Preoperative and postoperative CTA images, along with accompanying clinical data, were retrospectively retrieved from the CUMC database for an ATAA patient with STJ dilation and a control subject with no known aortic disease (Table 1). All data were de-identified in accordance with established protocols at CUMC and handled in accordance with HIPAA-compliant procedures throughout the study. CTA imaging was performed on a Siemens Biograph 128 scanner with intravenous contrast, gated to acquire during the diastolic phase of the cardiac cycle. The resulting images have a field of view (FOV) of 389 mm with a matrix size of 512 × 512, resulting in a pixel size of 0.79 mm and a slice thickness of 0.60 mm for the ATAA patient. The control subject has the same matrix size but an FOV of 220 mm, resulting in a pixel resolution of 0.43 mm and a slice thickness of 0.5 mm. Postoperative CTA imaging was performed on the ATAA patient approximately six months after the surgery. We note that the control patient’s CTA showed mild calcifications on the aortic valve, although the ascending aorta was normal. We assumed the valve is normal for the purpose of our analysis.

**Table 1.**
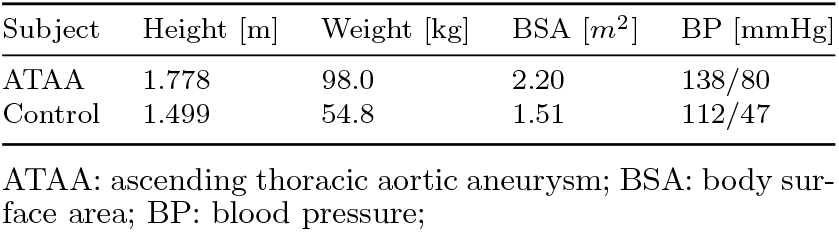
Clinical data for the two study subjects.

| Subject | Height [m] | Weight [kg] | BSA [$m^2$] | BP [mmHg] |
| --- | --- | --- | --- | --- |
| ATAA | 1.778 | 98.0 | 2.20 | 138/80 |
| Control | 1.499 | 54.8 | 1.51 | 112/47 |

ATAA: ascending thoracic aortic aneurysm; BSA: body surface area; BP: blood pressure;

### 2.2 Model Creation

Below, we discuss major steps in the computational model creation pipeline for CFD and FSI analysis (Fig. 1a-c). The pipeline is applied to the preoperative (preop) and postoperative (postop) images of the ATAA patient and to the subject with no ascending aortic dilation (control).

**Fig. 1.**
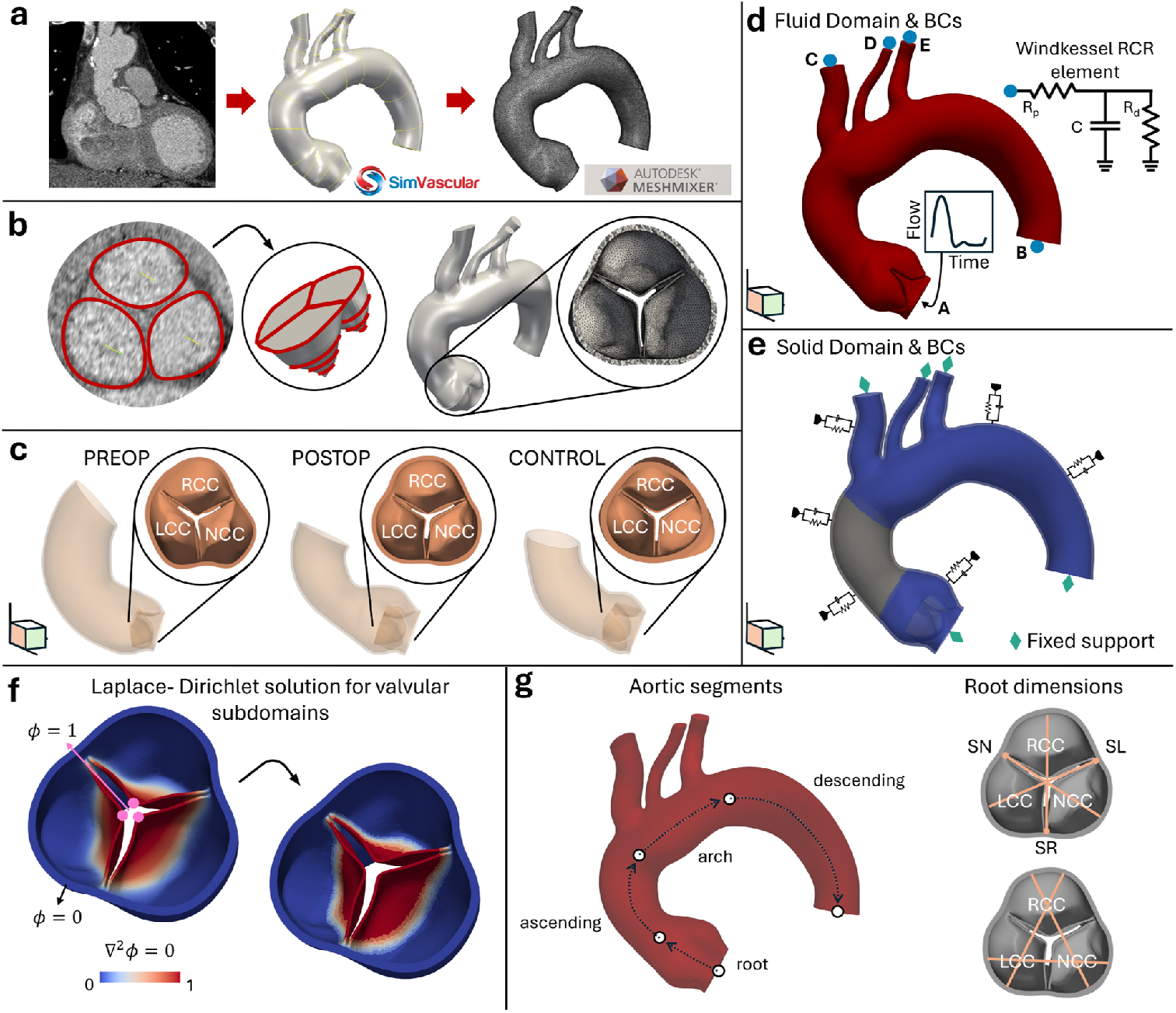
Overall workflow for creating and analyzing a patient-specific FEA model of the thoracic aorta with the aortic valve, including boundary condition configuration and material subdomains for CFD and FSI analyses. (a) Pipeline to extract aortic vessel lumen from a patient’s CTA images using SimVascular and MeshMixer (Autodesk, Inc.), including path planning, model lofting, and meshing [20, 21]. (b) Steps to segment a patient-specific aortic valve, fully integrated with the surrounding aortic vessel wall. The vessel wall is created by extruding the lumen’s outer surface by 1.5 mm using MeshMixer. (c) Cross-sectional views of the segmented aortic valve, viewed from the aortic side, for the three cases studied: preoperative (preop) and postoperative (postop) models of an ATAA patient with dilated STJ, and a control subject. (d) Fluid domain with relevant boundary conditions comprising a prescribed inflow profile at the inlet (A) and three-element Windkessel networks at each outlet (B-E). (e) Boundary conditions for the solid domain accounting for external tissue support on the outer wall, while the inflow and outflow annular walls are fixed. The highlighted gray subdomain corresponds to the reconstructed STJ graft, where the properties of Dacron material were applied to assess material sensitivity. (f) Laplacian-Dirichlet solution field (*ϕ*) used to construct valvular subdomains to allow a smooth transition in the material parameters between the leaflet belly and leaflet attachment edge. (g) Schematics for morphological analysis of (left) aorta partitioned into segments based on vessel centerline [22] and (right) aortic root, characterized using length measurements between (top) sinus to commissure and (bottom) commissure to commissure. FEA: finite element analysis; CFD: computational fluid dynamics; FSI: fluid-structure interaction; CTA: computed tomography angiography; ATAA: ascending thoracic aortic aneurysm; STJ: sino-tubular junction; RCC: right coronary cusp; LCC: left coronary cusp; NCC: non-coronary cusp.

#### 2.2.1 Segmentation and meshing

We segmented models of the thoracic aorta from the patient’s gated CTA images following Sim-Vascular’s pipeline [20], which involves path planning, model lofting, and meshing (Fig. 1a). Any surface irregularities after model lofting, especially near the aortic cusps and branch anastomoses, were smoothed in MeshMixer (Autodesk Inc.). For the FSI analysis, the arterial wall is created by extruding the lumen outer surface by ∼ 1.5 mm along the surface normal [21]. This thickness is in agreement with average values reported for adult aortic tissue [23]. Surface distortions due to intersections near the branch vessels after extrusion are resolved by local smoothing operations in MeshMixer (Autodesk, Inc.). Each model consists of the aortic root, ascending aorta, aortic arch with the three upper branches, and a small portion of the descending aorta (Fig. 1).

Before detailing our aortic valve segmentation pipeline, it is important to discuss the nuances of the inflow plane location and orientation. Most studies performing blood flow analysis of the ascending aorta truncate the model at the root-STJ interface, excluding the aortic cusps and coronary ostia [24–31]. Alternatively, the truncated plane is positioned near the base of the aortic valve, thereby capturing the cusps and coronary arteries, although a majority of these studies do not incorporate the valve leaflets [32–34]. In both these cases, the inflow plane is typically oriented so that its normal is aligned with the vessel centerline. However, small deviations in the inflow plane could affect the velocity patterns in the aortic root, which are of primary interest in this study. Moreover, as the current study focuses on performing both CFD and FSI analyses, it is crucial to ensure that the inflow plane is not only identical between the two models but also accurately captures the flow orientation and its impingement on the aortic valve. To account for these sensitivities in the current study, the inflow plane is aligned with the leaflet attachment plane (i.e., where all the leaflets simultaneously attach to or emerge from the aortic wall) at the base of the aorta, and is positioned a few mm away from the basal plane in the left ventricular outflow tract (LVOT). Our workflow for capturing this leaflet attachment plane also informs valve segmentation, as demonstrated in our prior work [35], and described next.

Briefly, a two-point linear path is created in SimVascular that is perpendicular to the plane of leaflet attachment. Adding more points to the path selection could result in a curvy spline-fitted path, which is intentionally avoided to maintain the orientation of the segmentation plane. This path selection is accomplished during path planning by repeatedly traversing along a potential path and adjusting the image cross-planes so that all the leaflets emerge from the tissue simultaneously. However, since CTA images are generally oriented in anatomical planes (axial, coronal, and sagittal), they do not automatically capture the leaflet attachment plane. Therefore, this plane adjustment is facilitated by SimVascular’s cross-plane rotation tool, which allows the independent rotation of each anatomical plane, thereby creating a multiplanar reconstructed view of the aortic valve. Our approach to aligning the valve segmentation path with the leaflet attachment plane normal enables the capture of inter-leaflet morphological differences, which have been increasingly studied recently in the context of normal and pathological aortic valve [35–37], and deviates from most models that assume leaflet symmetry [38–40].

Once the transvalvular path is determined, we then proceed to segment the leaflets, with the ultimate goal of integrating them into the aortic wall as a unified tissue (i.e., the solid domain). Each leaflet is segmented by manually tracing its ring-like contour at multiple longitudinal locations along the path [35], keeping the portion coinciding with the arterial wall external to the previously segmented aortic lumen boundary (Fig. 1b). The method of extending the leaflet beyond the lumen’s outer surface and then clipping the excess ensures no gaps are present between the two domains once they are joined to form a unified solid domain. The leaflets were extruded 0.6 mm normal to the segmented surface. Each leaflet was joined with the vessel wall using Boolean operations, and the resulting junctions were smoothed. The path planning, leaflet segmentation, and lofting are performed in SimVascular [20, 35], while model clipping, boolean operations, and smoothing are performed in MeshMixer (Autodesk, Inc.) [35].

Both the CFD-ready lumen and the FSI-capable solid model volumes are meshed inside SimVascular using TetGen with linear four-noded tetrahedral (TET4) mesh elements. For the CFD analyses, we incorporated boundary layer meshes to enhance near-wall resolution, thereby improving the accuracy in resolving velocity gradients and capturing relevant flow characteristics. The entire solid model maintains at least two elements across the tissue for both the arterial wall and valve leaflets. The inner surface of the solid model is reimported into SimVascular, and its faces are capped to create a lumen domain for the FSI analysis [21, 30, 41]. Because this workflow uses identical surface meshes at the solid domain’s inner wall and the fluid domain’s outer wall, a strong coupling is automatically achieved via node-matching to satisfy kinematic boundary conditions at the lumen-solid interface (Section 2.3.3). We generated meshes with a maximum global element edge size ranging from 1 mm to 1.3 mm, consistent with previously published mesh convergence studies using a similar geometry [21]. Across all six modeled cases, including preop, postop, and control for CFD and FSI, each final meshed geometry consisted of ∼2-3M TET4 elements.

#### 2.2.2 Initial valve geometry

Our starting point for creating the FEA model is the patient’s gated CTA images, acquired during diastole when the aortic valve is closed. As a result, the segmented valve is in its closed state, with leaflets nearly in contact with each other and assuming a concave profile. Small gaps yet remain due to the smoothing operations performed during segmentation. During opening, when a positive transvalvular pressure gradient is applied at the inflow in an FSI simulation, the leaflets change configuration from concave to convex. The traveling pressure wave creates a non-uniform pressure distribution on the leaflets, opening the leaflets’ belly regions while bringing the tips closer (often into contact), leading to severe mesh distortion and simulation blow-up. Therefore, instead of using the segmented valve configuration as the initial configuration, we apply a small positive pressure on the leaflets and perform a quasi-static simulation to slightly increase the leaflet separation (Fig. 1c). This gap also allows for creating a fluid mesh with sufficient density between the leaflets, which facilitates smooth deformation as the leaflets open without severe mesh distortion and voids the need for remeshing during the course of simulation.

#### 2.2.3 Solid domain separation

The final solid model from the segmentation pipeline (Section 2.2.1) is a unified tissue model that smoothly integrates the aortic vessel wall with the valve tissue. However, biomechanical experiments show substantial differences in the material properties between these tissues [42, 43], and separating these domains is not straightforward without manual intervention. To achieve this objectively, we solved a Laplace equation for a scalar field to smoothly diffuse across the entire tissue with a Dirichlet boundary condition applied at the center of each leaflet’s free edge. A threshold was then selected to split the domain into valve and arterial wall for the corresponding material property specification. To evaluate the sensitivity of the predicted hemodynamics to varying structural heterogeneity, we added a transitional domain between the leaflet’s belly and the attachment edge through which the material properties smoothly transition between the valve leaflets and the aortic wall (Fig. 1f). Further, an additional domain was created in the postop model to represent the clinically reconstructed STJ with a polymeric Dacron graft (Fig. 1e). An FSI simulation was then set up using the graft’s material properties in this reconstructed STJ domain, while the native tissue parameters were used in the rest of the aortic wall.

### 2.3 Governing Equations

Here, we provide a brief overview of the governing equations and boundary conditions for both CFD and FSI problems. Section 2.3.1 describes the fluid subproblem using the arbitrary Lagrangian-Eulerian (ALE) formulation for moving domain blood flow simulations, which is also extended to the CFD problem with rigid boundaries. Section 2.3.2 provides an overview of the solid subproblem and associated boundary conditions, while the interface conditions to enable FSI are discussed in Section 2.3.3.

#### 2.3.1 Fluid subproblem and boundary conditions

Blood flow through the ascending aorta is modeled as incompressible and Newtonian, governed by the Navier-Stokes equations, written in ALE coordinates for moving domains. With **v**_*f*_ being the fluid velocity (Eulerian) and 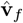 the velocity of the fluid domain, which deforms arbitrarily in response to the interfacial movement (Lagrangian), the governing momentum and mass conservation equations are given as [44–46],

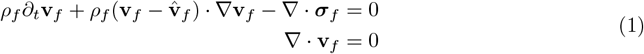

where *ρ*_*f*_ is the fluid density and ***σ***_*f*_ is the Cauchy stress tensor for a Newtonian fluid,

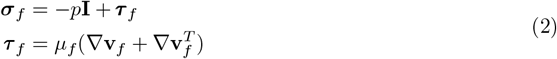

where *p* is fluid pressure, ***τ*** _*f*_ is the viscous stress, and *µ*_*f*_ is the dynamic viscosity of the fluid. In the current study, we set *ρ*_*f*_ to be 1.06 g/cm^3^ and *µ*_*f*_ as 0.04 Poise, which match commonly used parameters for blood [21, 32].

Dirichlet boundary conditions were applied at the inlet in the form of a generic unsteady blood flow profile (Fig. 1d). While the inlet profile remains the same for the preoperative and postoperative cases of the ATAA patient, when applied to the control subject, the profile was scaled to match the cardiac index (i.e., BSA-normalized cardiac output, Table 1), facilitating a reasonable comparison.

Each of the model outlets is connected to a three-element RCR-type Windkessel model (Fig. 1d), comprising a proximal resistance (*R*_*p*_) in series with a parallel combination of a capacitor (*C*) and a distal resistance (*R*_*d*_). The RCR parameters are tuned to match the cuff pressure measurements for the postoperative case (Table 1), and were retained for simulating the preoperative case as well. For the control, however, the RCR parameters were scaled using BSA as,

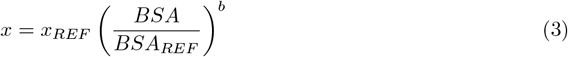

where *x* is a generic resistance (*R*_*p*_, *R*_*d*_) or capacitance (*C*) for each of the outlets (Fig. 1d). *x*_*REF*_ and *BSA*_*REF*_ represent the reference values associated with the postoperative case. The exponent *b* depends on whether *x* represents a resistor or a capacitor, as well as whether the outlet is connected to upper or lower body circulation [47, 48]. For resistance scaling, *b* = − 0.3 for upper branch vessels, while *b* = − 0.9 for the main arterial outlet in the descending aorta. For capacitance scaling, *b* = 0.4 for upper body branch outlets, while *b* = 1.2 in the descending aorta [47, 48].

Further, as part of the ALE formulation, the fluid domain is smoothly deformed by solving an element-Jacobian-stiffened linear elastostatics problem for the fluid mesh velocity 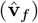 and displacements (**û**_*f*_), augmented by Dirichlet boundary conditions at the interface to satisfy kinematic equilibrium between the fluid and solid subdomains (Section 2.3.3) [44–46].

The governing equations for the CFD problem are the same as the fluid subproblem for the FSI (Eq. 1), excluding the domain velocity 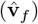 in the nonlinear convection term. We applied the same boundary conditions at the inlet and outlets of the fluid domain for the CFD cases that were used in the FSI simulations. However, a no-slip, no-penetration boundary condition is applied at the lumen’s outer surface, resembling a rigid, stationary wall.

#### 2.3.2 Solid subproblem and boundary conditions

The solid domain constructed from medical images comprises the aortic wall and valve as two separate subdomains, with a transitional domain in some cases to allow smooth variation in material properties (Section 2.2.3). Each of these subdomains is modeled as an isotropic and homogenous material undergoing finite deformations, governed by finite strain elastodynamics, and whose mechanical response is characterized as nonlinear and hyperelastic.

Briefly reviewing the concepts of finite strain elasticity [49], we define a deformation map, 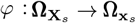 such that **x**_*s*_ = *φ* (**X**_*s*_, *t*), where **x**_*s*_ is the current position of a material point at a time *t* that was originally at **X**_*s*_ in the reference configuration. Here, we define 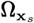 as the current or spatial configuration of the solid domain and hypothesize the existence of a reference unloaded configuration, 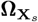. The displacement, **u**_*s*_, of a material particle and the deformation gradient tensors are given by,

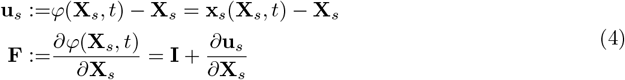

where, **F** is the deformation gradient, *J* := det (**F**) is its Jacobian determinant, and **C** = **F**^*T*^ **F** is the right Cauchy-Green deformation tensor. In the absence of any body forces, the governing elastodynamics equation is given in the reference configuration as,

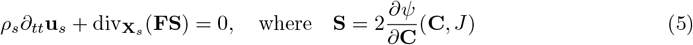

is the second Piola-Kirchhoff stress tensor, *ρ*_*s*_ is the density of the solid domain, 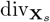 is the divergence operator in the reference configuration, and *ψ* is the strain energy density function for a hyperelastic material. In this work, we model the isochoric (i.e., volume-preserving) deformations of both arterial and valvular tissue using the neo-Hookean constitutive model, while the volumetric (dilatational or volume-changing) component of the strain energy is modeled using the Simo-Taylor 1991 (ST’91) constitutive model [50, 51]. The combined strain energy density function is given as [21],

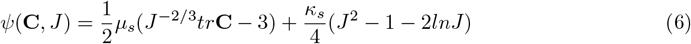

where *µ*_*s*_ and *κ*_*s*_ denote the shear and bulk modulus respectively, calculated from the material elastic modulus (*E*_*s*_) and Poisson ratio (*ν*_*s*_),

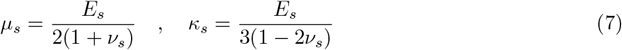

In the current work, *E*_*s*_ was set to 1 MPa for the arterial wall, and 0.4 MPa for the leaflets, while *ν*_*s*_ = 0.45 for the entire tissue based on previously published studies [52, 53].

Zero-displacement Dirichlet boundary conditions were applied at the solid domain annuli. A Robin-type boundary condition defined as,

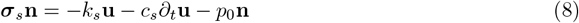

was applied to the outer wall surface to account for external tissue support and limit non-physiological movement such as twisting (Fig. 1e), which has been applied in other studies of the aorta, as well as other cardiac mechanics models [21, 30, 46, 54–57]. The parameters *k*_*s*_ and *c*_*s*_ model the elastic and viscoelastic response of the external tissue, while *p*_0_ represents the intrathoracic pressure [54]. For our purposes, we prescribed a *k*_*s*_ value of 10^6^ dyn*/*cm^3^ and *c*_*s*_ = *p*_0_ = 0, consistent with previously published literature [21].

#### 2.3.3 Interface conditions

In our ALE formulation for FSI (Section 2.3.1), the fluid and solid subdomains are strongly coupled at the interface, satisfying kinematic and dynamic boundary conditions. A kinematic equilibrium at the interface ensures a balance of nodal velocities and displacements (i.e., 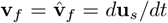, and **û**_*f*_ = **u**_*s*_). On the other hand, the interfacial stress balance ensures dynamic equilibrium (i.e., ***σ***_*f*_ = ***σ***_*s*_), where ***σ***_*s*_ is the Cauchy stress in the solid domain. As a result of our fluid and solid mesh creation workflow leading to a node matching between the two subdomains at the interface (Section 2.2.1), the kinematic boundary conditions are strongly enforced, while the dynamic boundary conditions are matched to the tolerances up to which the weak form residuals are satisfied.

### 2.4 Simulation setup

All the simulations in this study are performed using our in-house multiphysics finite element solver, adapted from the open-source finite element solver, *svFSI* ^2^ [58], which has been previously validated and employed to model a variety of cardiovascular biomechanics applications, including FSI modeling in aortic dissection and aneurysms [21, 59, 60], blood flow in coronaries [61, 62], cardiac mechanics [46, 56, 57, 63, 64], electrophysiology [65], hemodynamics in developing ventricles [45, 66], and multiphysics applications [67], including vascular growth and remodeling [68, 69]. We employed stabilized finite elements to address numerical instabilities caused by strong convection and circumvent the *inf-sup* conditions associated with mixed finite element formulation [44]. Time integration was performed using the implicit, second-order accurate, generalized-*α* method [70, 71]. The resulting nonlinear system of equations are solved using the Newton-Raphson method, embedded within a multi-step predictor-corrector method [44, 45, 72].

For the CFD problem, within each nonlinear solver iteration, we employed the bi-partitioned method (BIPN) to efficiently solve the system of linear equations [73]. For the FSI problem, the fluid and solid system of equations are monolithically coupled, while a loose coupling is adopted for solving the deformation of the fluid domain [74]. Further, within each block-wise nonlinear iteration, we employed the iterative solver generalized minimal residual (GMRES) [75] linear solver with diagonal preconditioning to solve the linearized ALE system of equations (Eqs. (1, 5)), whereas the Conjugate Gradient (CG) method is used to solve the linearized system of equations governing the fluid mesh motion. To handle the 0D/3D coupling between the 3D fluid domain and the RCR boundary conditions, a resistance-based preconditioning is employed to accelerate the convergence of the linear solver [63, 76, 77]. The nonlinear solver tolerance was set to 10^−4^ while the linear solver tolerance was set to 10^−6^ for all cases.

The ATAA patient and the control subject were modeled using the same heart rate, 75 BPM, with a time step size of 0.0008s for the CFD simulations, yielding 1000 time steps per cardiac cycle. The time step size for FSI simulations was lowered by half to avoid numerical instabilities. The CFD simulations were run for six complete cardiac cycles to reach a limit cycle, while the FSI cases were run only through the leaflet opening phase (ventricular systole). Each CFD simulation typically takes approximately 1.3 hours for one cardiac cycle using 128 AMD-EPYC-7742 CPU cores on the Expanse supercomputing cluster at UC San Diego, while the FSI simulation takes approximately 4 hours on the same cluster to simulate valve opening. A summary of all the simulation parameters and solver settings is provided in Table 2.

**Table 2.**
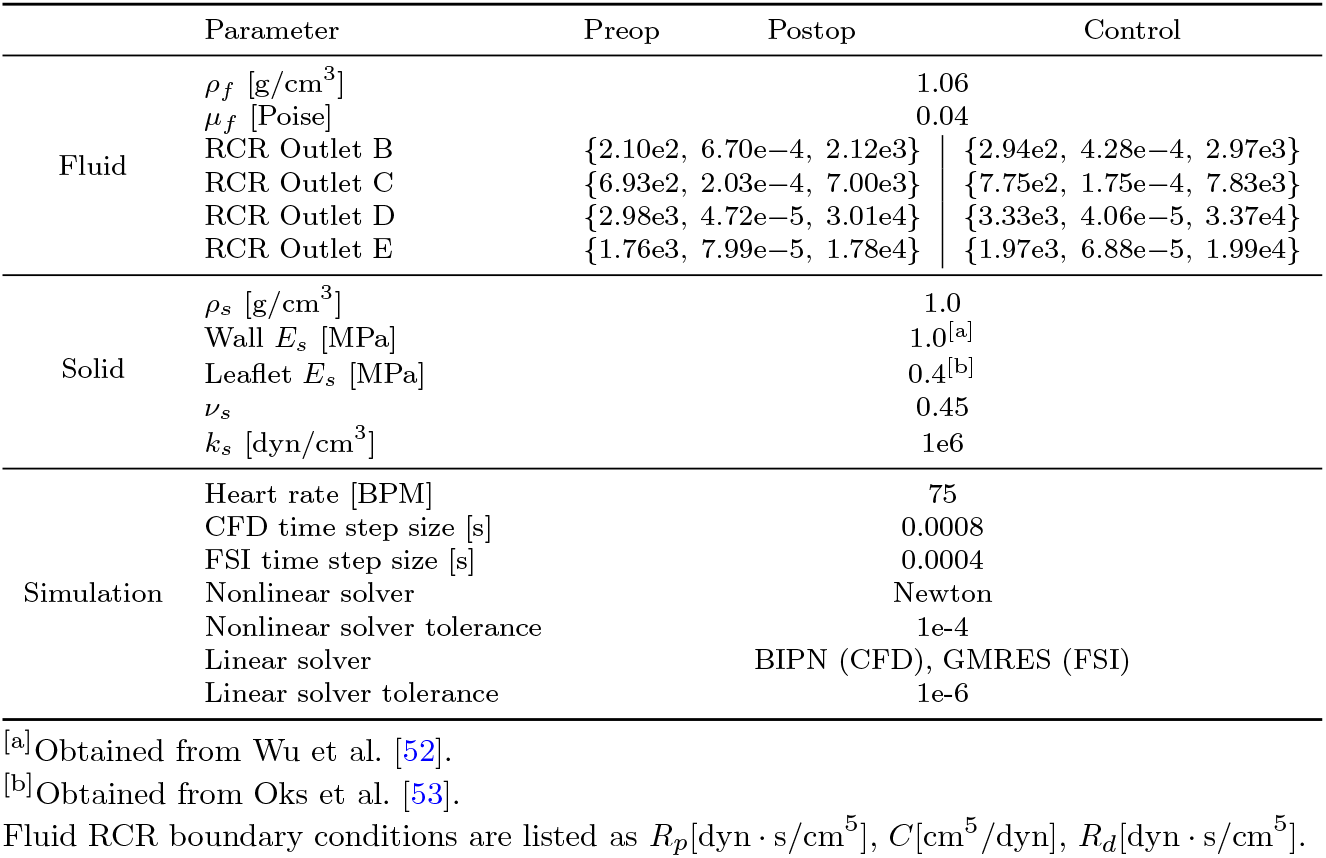
All parameters used for simulations.

### 2.5 Evaluation Metrics

In this study, we performed morphological and hemodynamic analyses of the thoracic aorta for all cases considered (i.e., preop, postop, and control). Morphological analysis includes vessel diameter, aortic root dimensions, and leaflet areas. Hemodynamic analysis includes stream traces, mean velocities and flow rates, pressure and shear forces, as well as the kinetic energy and viscous dissipation. Additionally, we also analyzed leaflet opening characteristics in the FSI simulation and the sensitivity of the tissue displacements and strains to material properties.

We extracted the vessel diameter along the length of the thoracic aorta, partitioned into segments that included the root, ascending aorta, aortic arch, and the descending aorta (Fig. 1g), based on the classification provided by Hager et al. [22]. The root region was defined as the region from the inlet plane to the STJ. The ascending aortic portion was determined to be the vessel volume leading up to the brachiocephalic trunk from the STJ. The aortic arch was chosen as the region spanning the three major upper branches, and the rest of the vessel was labeled as the descending aorta [22]. Briefly, the vessel centerline is extracted from the segmentation in SimVascular as a series of points with a typical resolution of 0.35 mm. The point coordinates of the inner surface of the vessel wall are transformed from Cartesian to cylindrical coordinates at each node with respect to the vessel centerline, and each surface node is mapped to its nearest centerline point. The vessel length is uniformly sampled into segments, with each segment spanning ten centerline points. The average diameter is then determined for each segment and plotted as a function of the vessel centerline coordinate.

While the notion of ‘diameter’ applies to a circular cross-section and is valid for the bulk of the aorta, it is not straightforward to define a ‘diameter’ for the aortic root, which has a cloverleaf anatomy with three sinuses [78]. Clinically, the size of the aortic root is characterized using two types of lengths: length along a sinus to the opposite commissure and length along a sinus to the adjacent sinus (Fig. 1g) [79, 80]. Recent advances suggest adopting the diameter of a circumscribing circle around the aortic root to closely approximate wall stress equilibrium with blood pressure based on the Laplace law, although this is yet to be adopted into clinical practice [78]. In this work, we measured distances from a sinus to the commissure for each of the coronary cusps (*L*_*SR*_, *L*_*SL*_, *L*_*SN*_) and distances between sinuses (*L*_*LR*_, *L*_*RN*_, *L*_*NL*_). The sinus-to-commissure measurement is obtained by computing the distance between each labeled commissure (SR, SL, and SN in Fig. 1g) and the opposite sinus at the point of peak convexity. The sinus-to-sinus measurement is obtained by measuring the maximum distance between adjacent sinuses (Fig. 1g). All measurements were conducted on the inner wall within the leaflet attachment plane. Finally, from the final tissue model with a fully integrated aortic valve, the leaflet area was calculated by averaging the areas of the aortic and ventricular sides of each individual leaflet.

In addition to the morphological analysis, hemodynamic metrics were used as comparison points between the three cases. These include domain-averaged velocities in the aortic root as well as flow rates and area-averaged pressures at inlet and outlet faces. We also computed shear metrics such as the time-averaged wall shear stress (TAWSS) and oscillatory shear index (OSI), defined as,

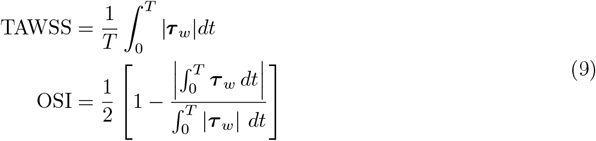

where 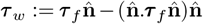 is the tangential wall shear stress vector acting on a surface with a unit normal 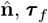 is the fluid viscous stress (Eq. (2)), and T is the duration of interest. Additionally, we calculated the volume-averaged kinetic energy density 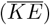 and viscous dissipation 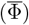, defined as,

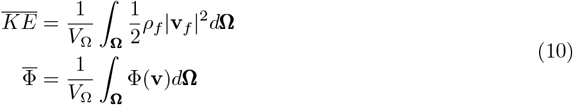

where *V*_Ω_ is volume of the domain of interest Ω, Φ(**v**) = 2*µ*_*f*_ ***ε*** : ***ε*** for a Newtonian fluid, and 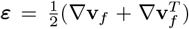. All these metrics are evaluated from the final converged cycle after a limit cycle has been established, and have been widely used in computational modeling of blood flow [21, 27, 41, 45], as they can help visualize certain blood flow patterns and behaviors that are potential indicators of abnormal hemodynamics.

## 3 Results

### 3.1 Morphological analysis

The centerline diameter analysis confirms that the ATAA patient has a dilated ascending aorta, with diameters reaching a maximum of over 4.5 cm around the mid-ascending aorta (preop in Fig. 2a). Post surgery, the STJ region shows a marked reduction in diameter, reaching the levels of the descending aorta for this patient (postop in Fig. 2a). Although the segmental diameters for the control subject are much lower than the ATAA patient post surgery (control vs. postop in Fig. 2a), these are still within ±2 standard deviations around population averages in adults with a healthy aorta [22].

**Fig. 2.**
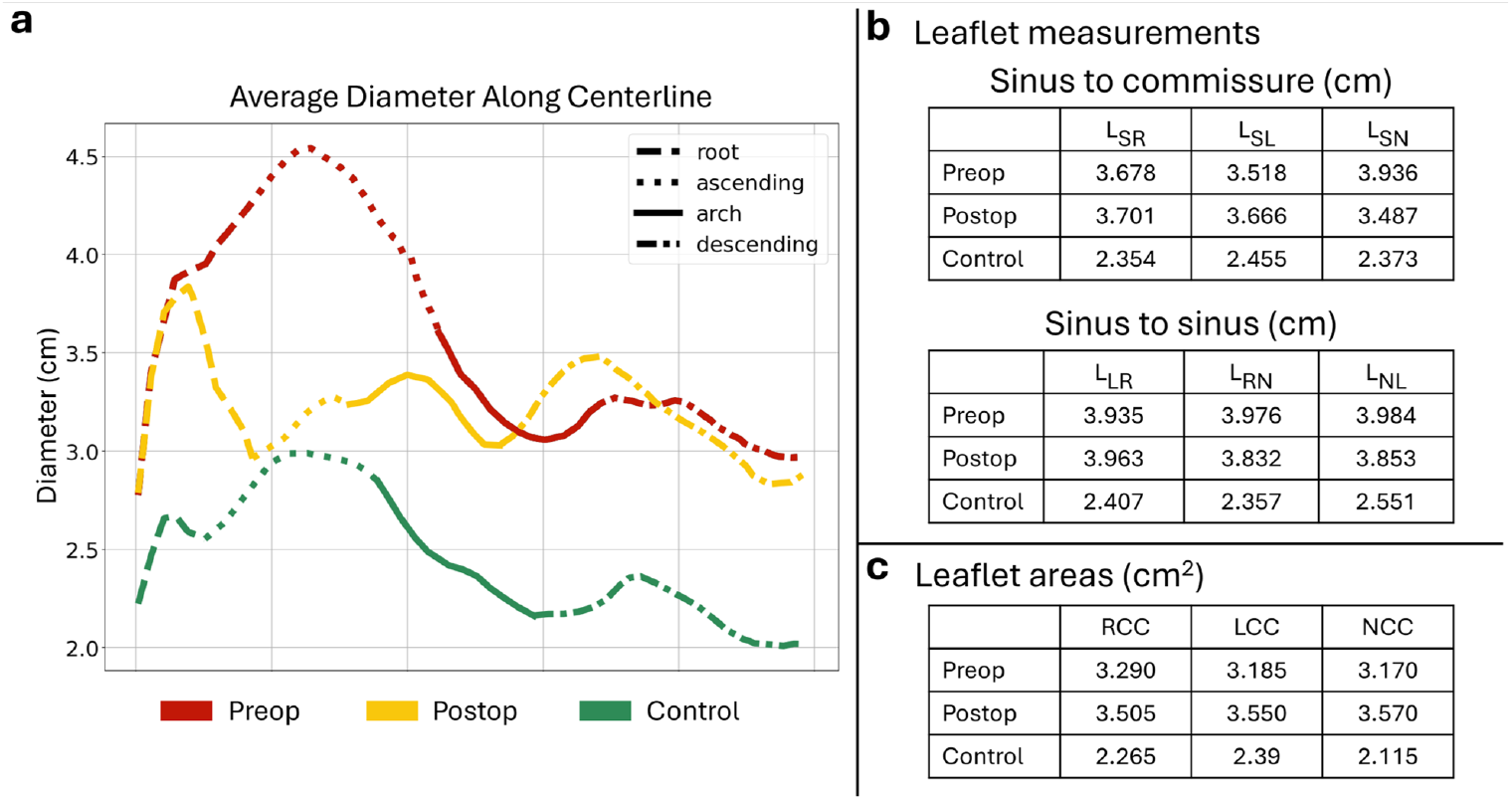
Morphological analysis of the thoracic aorta, aortic root, and aortic valve leaflets for the ATAA patient (preop, postop) and the control subject. (a) Section-wise mean diameter along the vessel centerline for all three study cases. Highlighted sections include the root, ascending aorta, aortic arch, and descending aorta, as defined in Fig. 1g. (b) Measurements of the aortic root made along the (top) sinus to commissure and (bottom) sinus to sinus, as defined in Fig. 1g. (c) Surface area of each leaflet. ATAA: ascending thoracic aortic aneurysm; RCC: right coronary cusp; LCC: left coronary cusp; NCC: non-coronary cusp.

Mean sinus-to-commissure lengths for the preop, postop, and control are 3.71 ± 0.21 cm, 3.62 ± 0.11 cm, and 2.39 ± 0.05 cm, respectively (Fig. 2b, top). Mean sinus-to-sinus measurements for each of these cases are 3.97 ± 0.03 cm, 3.88 ± 0.07 cm, and 2.44 ± 0.10 cm, respectively (Fig. 2b, bottom). Therefore, the characteristic size of the aortic root differs by nearly 7% for the ATAA patient when using the sinus-to-commissure measurement compared to sinus-to-sinus, whereas it only differs by 2% for the control subject. Likewise, the mean leaflet area for the preop, postop, and control cases is given by 3.22 ± 0.07 cm^2^, 3.54 ± 0.03 cm^2^, and 2.26 ± 0.14 cm^2^, respectively (Fig. 2c).

### 3.2 CFD Analysis

Computed flow rate and pressure waveforms at the domain boundaries show strong physiological relevance in both shape and magnitude [81], indicating a reasonable simulation setup (Fig. 3). Waveforms for the preoperative and postoperative cases show negligible differences, with a total cardiac output of 5.4 liters/min and a min/max aortic pressure of 131/80 mmHg at the vessel inlet, which is within 6% of the clinical measurement (Table 1). The cardiac index-matched inflow profile for the control subject (Section 2.3.1) leads to a reduced cardiac output of 3.7 liters/min and min/max aortic pressures as 119/70 mmHg (Fig. 3).

**Fig. 3.**
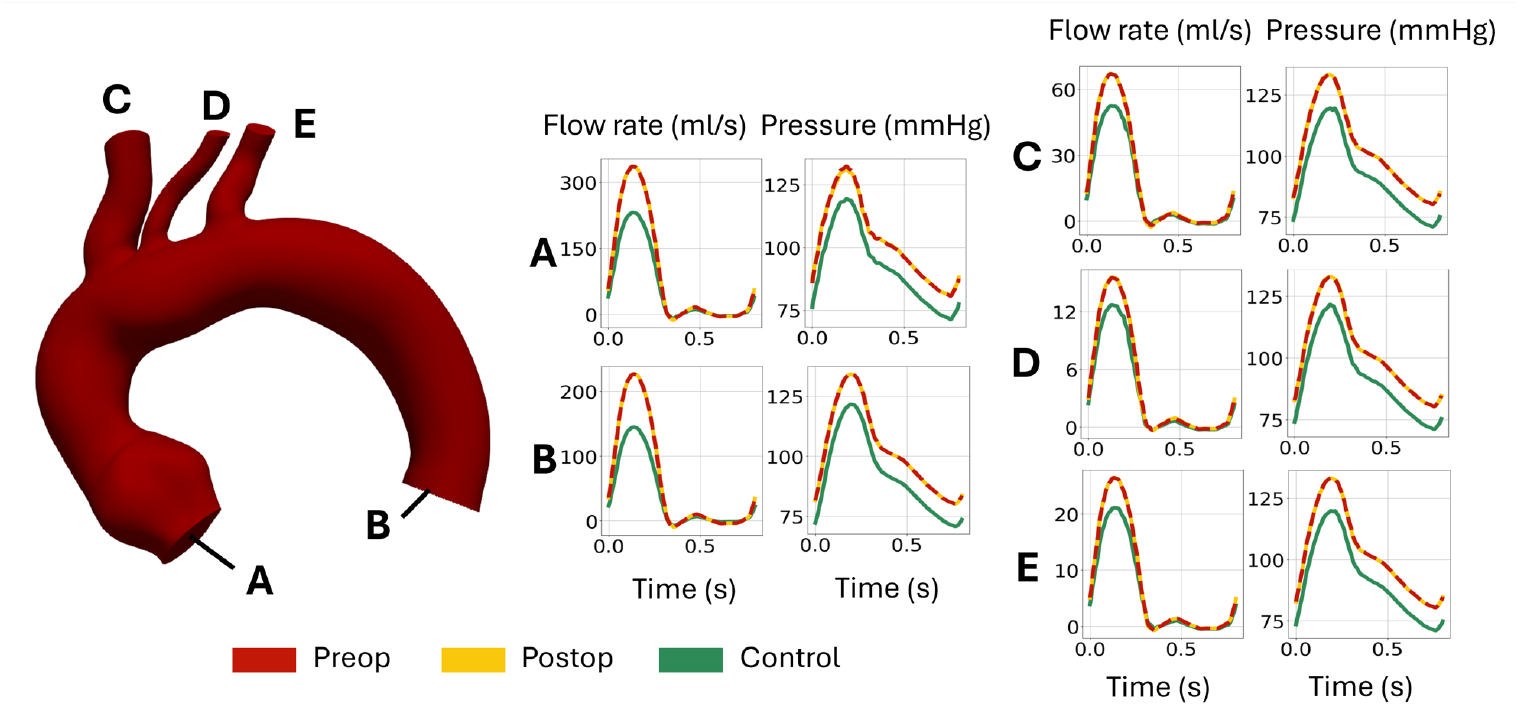
Comparison of CFD-derived flow rates (ml/s) and average pressures (mmHg) at the aorta inlet (A) and outlets (B-E) between preop, postop, and control.

The flow characteristics for all three models were found to be similar during the deceleration portion of the cardiac cycle; however, marked differences were observed between the preoperative and postoperative cases near peak systole (Fig. 4, preop vs. postop). Flow patterns in the aortic root for the postoperative case resemble those of the control subject, as evidenced by the presence of vortical structures in the sinuses and a more centrally aligned jet that retains its intensity throughout the ascending aorta (Fig. 4a). A separation bubble, characteristic of flow in curved tubes [82], is formed distal to the STJ in the ascending aorta (Fig. 4a). In contrast, the preoperative case shows a skewed jet that is swept against the outer curvature of the ascending aorta due to a large vortex in the mid-ascending aorta. Further, the root sinuses do not exhibit any prominent vortices, while the inflow jet quickly loses its intensity at the level of STJ (Fig. 4a, preop).

**Fig. 4.**
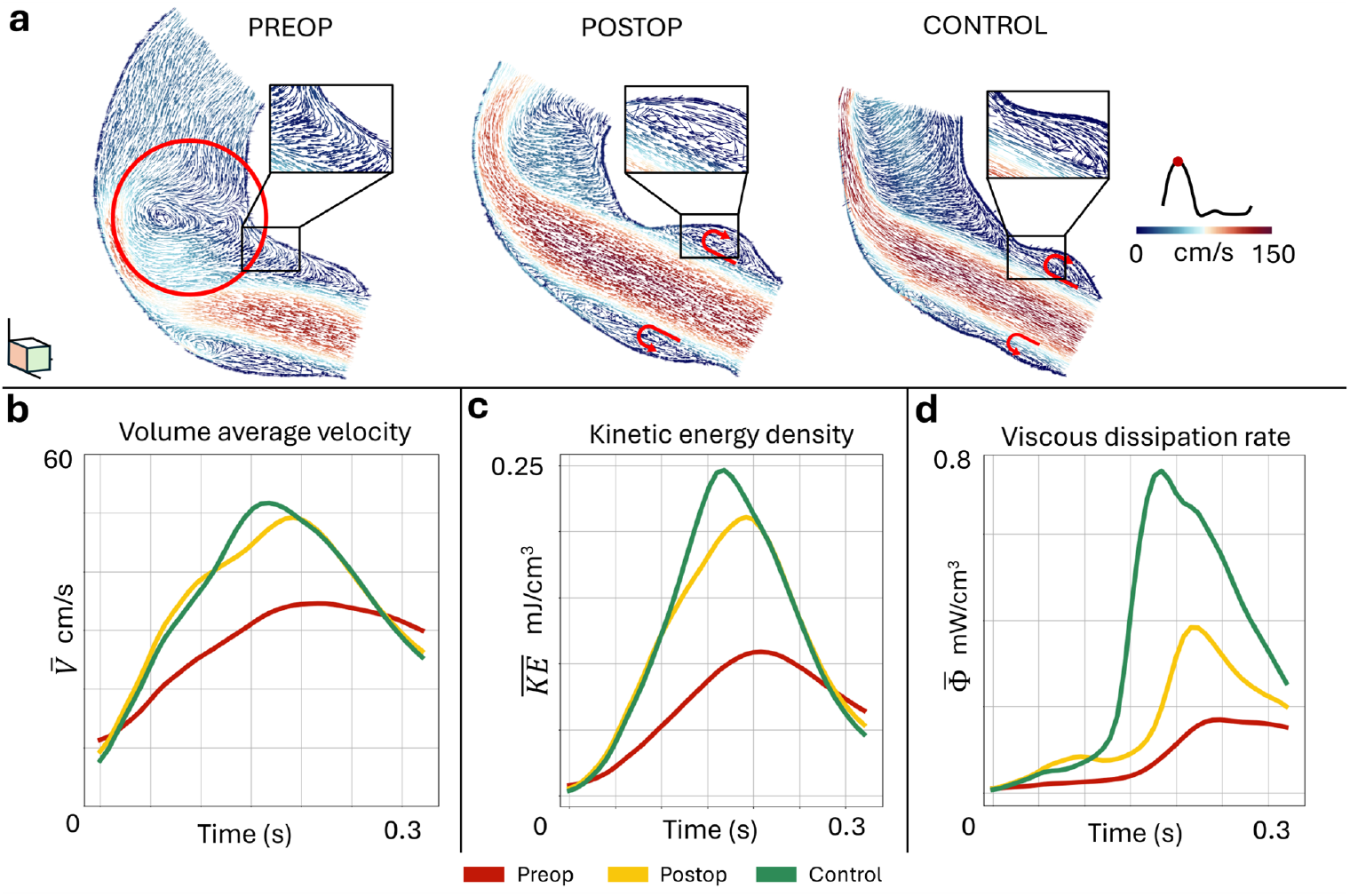
(a) Vector plots of the CFD-derived velocity field along a mid-plane capturing the root and ascending portions of the aorta near peak systole for all three cases. Insets highlight vortical structures near the root sinuses in the postop and control cases, which are absent in the preop case. (b-d) Comparison of volume-averaged velocity 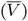, kinetic energy density 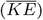, and viscous dissipation rate 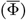 between preop, postop, and control during systole. The region of interest for volume-averaging encompasses the root and ascending portions of the aorta.

A comparison of the volume average velocity further demonstrates that the postoperative case more closely matches the control with its higher average velocities compared to the preoperative case (Fig. 4b). Similarly, the control case also exhibits the highest levels of kinetic energy density and rate of viscous dissipation in the aortic root, followed by the postoperative and the preoperative cases (Figs. 4c, d). We also note a sharp increase in the viscous dissipation rate for the control and postoperative models during early systole when the inflow jet impinges on the ascending aortic wall, compared to a relatively shallow increase for the preoperative model (Fig. 4d).

Hemodynamic shear profiles, including TAWSS and OSI, qualitatively show a close agreement between the postop and control cases, although the control exhibits markedly higher TAWSS (Fig. 5). The preoperative case, however, differs in terms of both magnitude and the region of high TAWSS and OSI. The control and postoperative cases display regions of high TAWSS distal to the STJ near the first superior branch (Figs. 5b,c), whereas the preoperative case is characterized by high shear occurring lower along the ascending aorta, proximal to the STJ (Figs. 5a). Further, both postoperative and control cases show a pocket of very low TAWSS in the mid-ascending aorta, which is lacking in the preoperative case. When examining the OSI in the root and ascending aortic regions (Fig. 5d-f), we see higher values in the root region for the postoperative and control cases, compared to the preoperative case. These shear characteristics are in agreement with observed jet profiles in the ascending aorta and vortical flow in the sinuses of the aortic root (Fig. 4a).

**Fig. 5.**
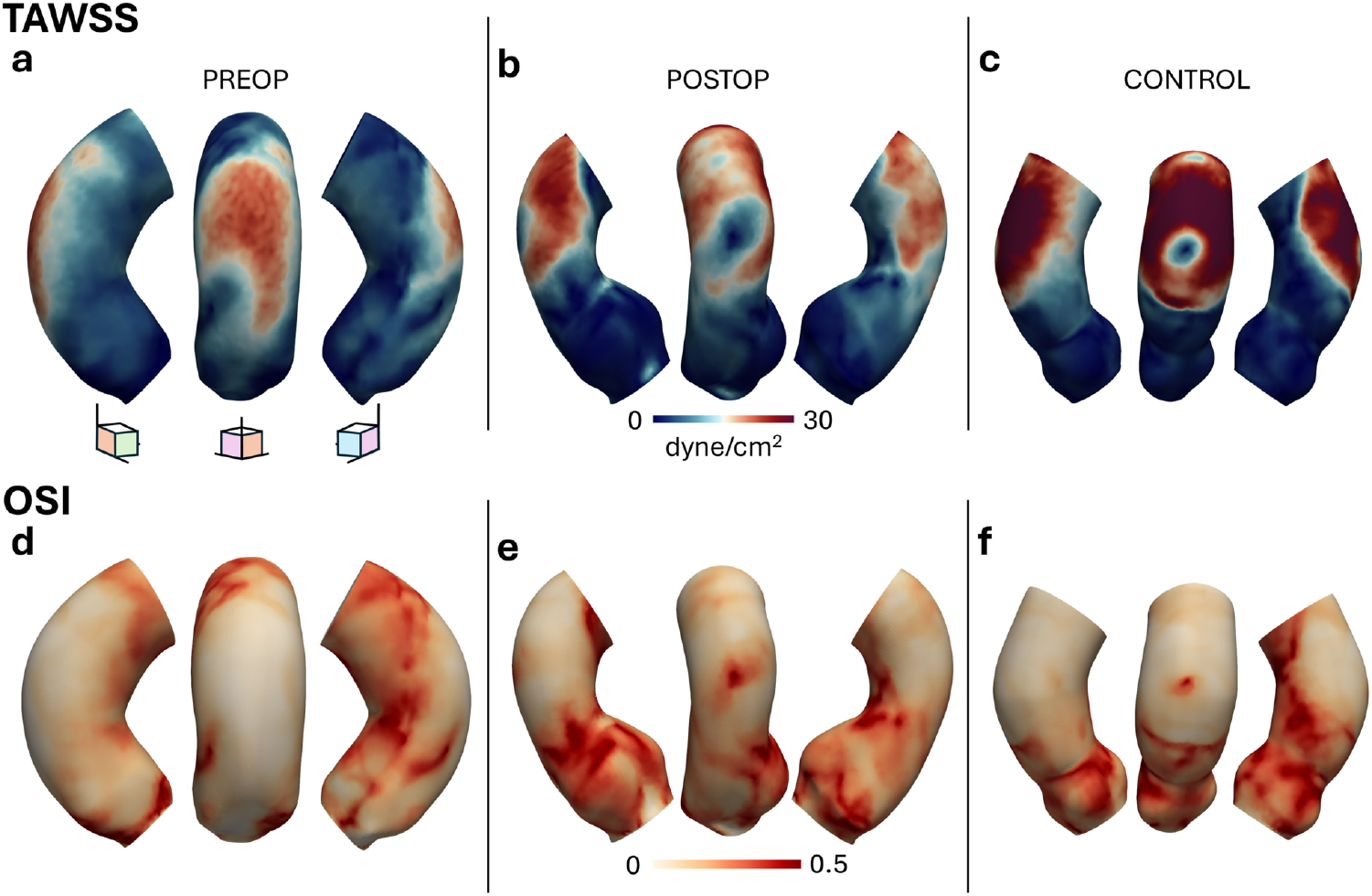
Comparison of CFD-derived shear profiles, including (top row) TAWSS and (bottom row) OSI, between preop, postop, and control cases, along three different views. TAWSS: time-averaged wall shear stress; OSI: oscillatory shear index.

### 3.3 FSI Analysis in Comparison to CFD

The flow patterns from FSI simulations, accounting for the deformation of the surrounding vessel under external tissue support and the aortic valve, show qualitative similarities with those of CFD-based flow features in the root and the ascending aorta (Fig. 6a). In particular, the control and postoperative cases are characterized by coherent sinus vortices, and a centrally positioned jet that impinges further along the ascending aorta near the first superior branch while maintaining its intensity throughout (Fig. 6a, control vs. postop). The relative size of the separated vortical flow in the ascending aorta with respect to the vessel diameter remains consistent between the two cases. In contrast, the preoperative case is characterized by weak sinus vortices, similar to the CFD result, although the leaflets align the inflow jet slightly more toward the center (Fig. 6a, preop). Furthermore, a large separated flow sweeps the inflow jet past the outer curvature, preserving the inflow kinetic energy throughout the length of the ascending aorta, unlike in the CFD case, where the inflow jet quickly dissipates near the STJ in the absence of the valve (compare preop in Figs. 4a and 6a). On the other hand, while the jet retains its magnitude throughout the ascending aorta for the postop case, the jet loses its intensity with a small secondary separation zone toward the end of the ascending aorta in the preop case (Fig. 6a, preop vs. postop).

**Fig. 6.**
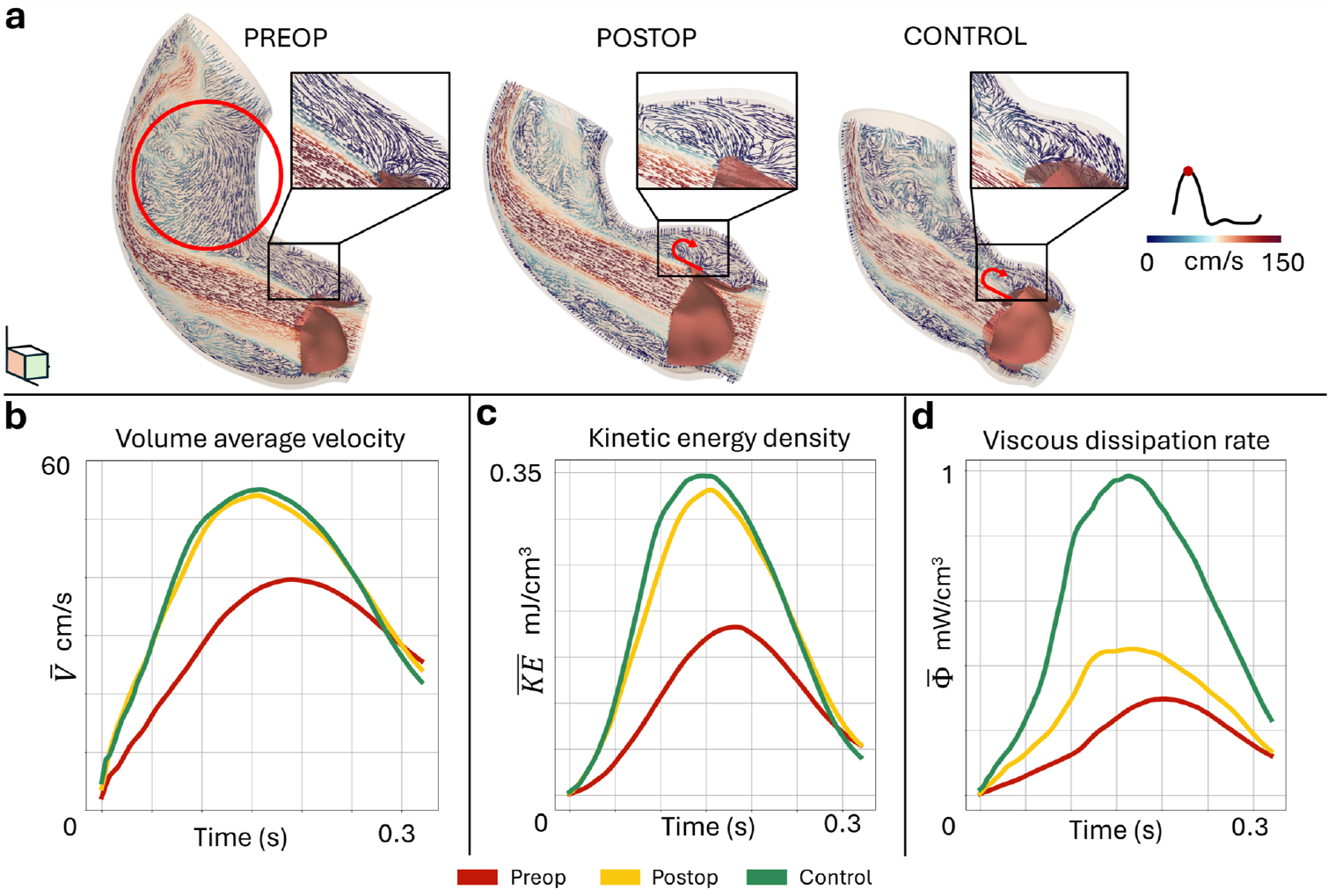
(a) Vector plots of the FSI-derived velocity field along a mid-plane capturing the root and ascending portions of the aorta near peak systole for all three cases. Insets highlight vortical structures near the root sinuses in the postop and control cases, which are absent in the preop case. (b-d) Comparison of volume-averaged velocity 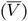, kinetic energy density 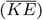, and viscous dissipation rate 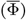 between preop, postop, and control during systole. The region of interest for volume-averaging encompasses the root and ascending portions of the aorta.

Quantitatively, we observed an overall increase in jet velocities, kinetic energy, and viscous dissipation rate for the FSI simulations compared to the CFD predictions, consistent with the valve dynamics that result in a narrower and temporally changing orifice area (Fig. 6b-d). Further, the general trends in these hemodynamic metrics between preoperative and postoperative cases remain consistent with the CFD, with increased magnitudes after surgery. However, including the aortic valve in the preoperative model resulted in a substantial increase in the peak values of volume-averaged velocities (↑ 15 %), kinetic energy density (↑ 68 %), and viscous dissipation rate (↑ 75 %), compared to the no-valve CFD model. Upon closer examination of the viscous dissipation rate, we observed a more gradual increase in the FSI simulations compared to a steep increase in the CFD data for control and postoperative cases during early to mid-systole (Figs. 4d and 6d).

Additionally, the TAWSS exhibits slight differences between CFD and FSI (Fig. 7a-c). While the location of high TAWSS is consistently lower along the ascending portion of the aorta for the preoperative case compared to the others in both modeling approaches, the magnitudes are generally elevated. When examining the OSI, we see a starker contrast between the regions of high and low OSI for each case compared to the CFD counterparts (Fig. 7d-f). However, caution must be exercised when comparing these shear indices against CFD data, due to the differences in the integrated time interval, which spans only the valve opening period in FSI, compared to the full cardiac cycle in CFD.

**Fig. 7.**
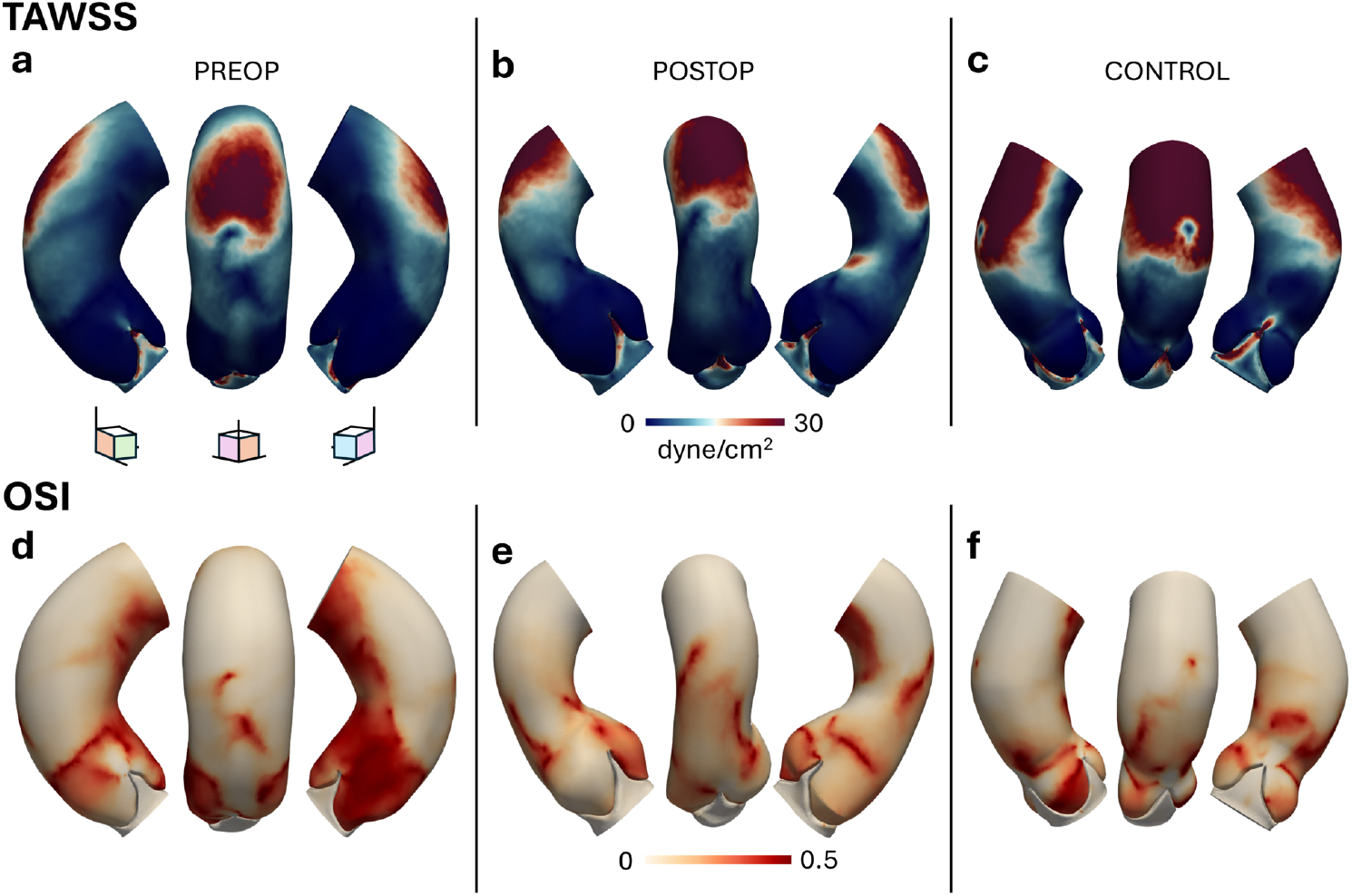
Comparison of FSI-derived shear profiles, including (top row) TAWSS and (bottom row) OSI, between preop, postop, and control cases, along three different views. Unlike the CFD case in Fig. 5, TAWSS and OSI for the FSI analysis are computed only during the aortic valve opening phase. TAWSS: time-averaged wall shear stress; OSI: oscillatory shear index.

To assess the leaflet dynamics during valve opening, we measured the peak geometric orifice area (GOA_max_) and systolic excursion of the leaflets (LE_sys_) for all three cases (Fig. 8). GOA_max_ for the preoperative and postoperative cases differ by 0.5%, with the preoperative case marginally higher than the postoperative model (Fig. 8a). However, the control shows a much lower GOA_max_ (2.02 cm^2^ for control vs. 3.178 cm^2^ for postop), attributed to the overall lower dimensions of the control subject’s aorta compared to the ATAA patient (Table 1). When normalized by BSA, however, the indexed GOA (GOAi) shows approximately 8% difference between the control and the ATAA patient, suggesting a reasonably normal opening dynamics of the aortic valve between the two subjects (Fig. 8a).

**Fig. 8.**
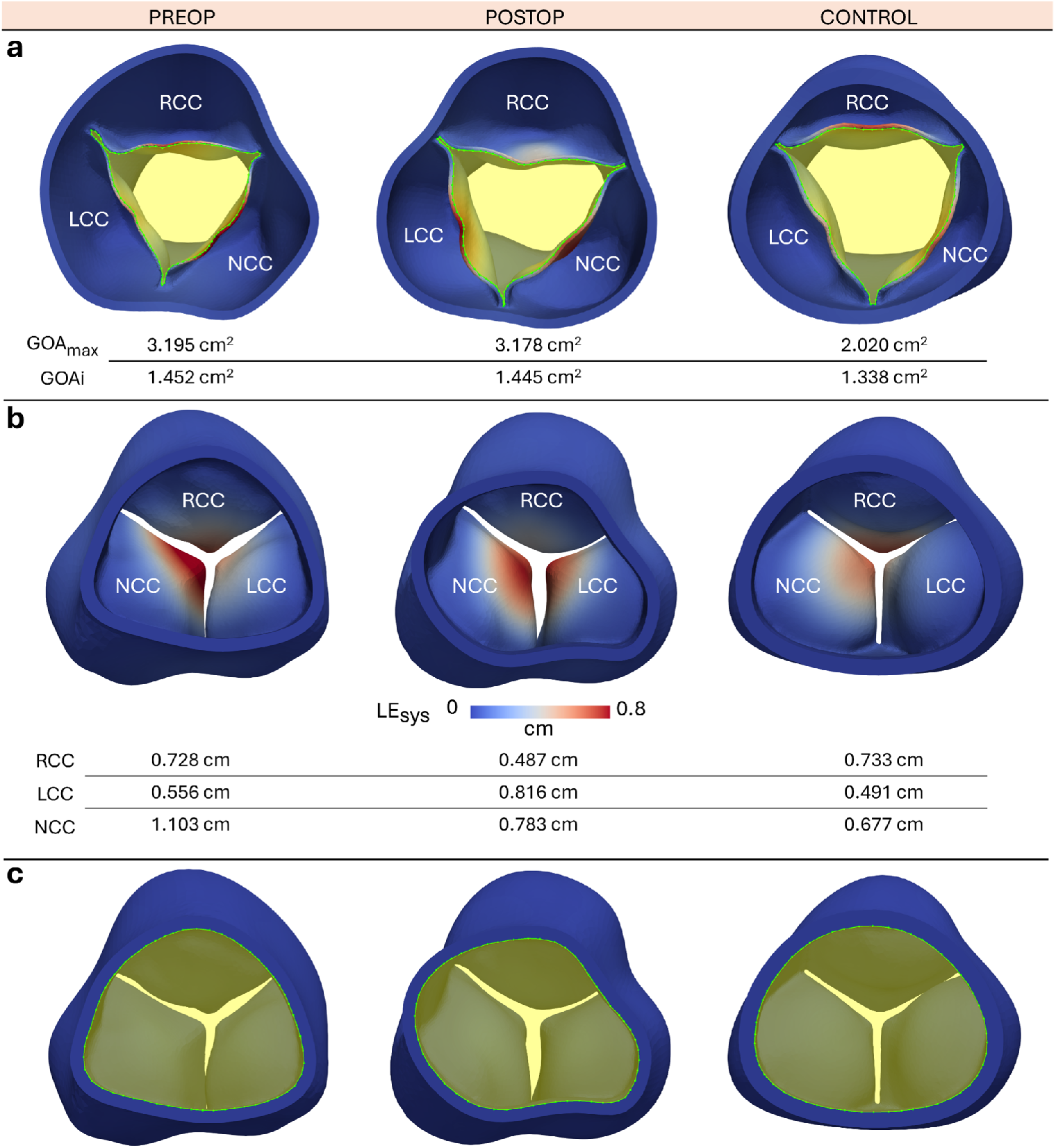
Analysis of the aortic valve opening dynamics for preop, postop, and control cases. (a) Cross-sectional view of the aortic valve at its maximal opening phase for each case. Both absolute GOA_max_ and indexed GOA (GOAi) are highlighted. (b) Maximum displacement of each leaflet, used as a surrogate for its systolic excursion LE_sys_. (c) Shape of the inlet lumen in the LVOT viewed from the ventricular side for each case. GOA: geometric orifice area; RCC: right coronary cusp; LCC: left coronary cusp; NCC: non-coronary cusp; LVOT: left ventricular outflow tract.

Systolic leaflet excursion (LE_sys_) analysis, measured as the maximum displacement of the individual leaflet from its diastolic configuration [35], indicates higher LE_sys_ for RCC and NCC than LCC in the control subject (Fig. 8b), which aligns with our previous observations in a cohort of twenty normal subjects [35]. However, the ATAA patient exhibits noticeable differences in the deformation of individual leaflets, although the overall orifice area was similar between the preoperative and postoperative cases and differed by a reasonable 8% compared to the control subject (Fig. 8). In particular, before surgery, NCC had the maximum LE_sys_, followed by RCC and LCC. Subsequent to the intervention, however, LCC has the maximum LE_sys_, followed by NCC and RCC. We also noted discrepancies in the LVOT profile at the base of the aorta in the ATAA patient, with an irregular post-surgery profile compared to the more rounded pre-surgery profile that also aligns with that of the control subject (Fig. 8c).

## 4 Discussion

In this retrospective computational study, we evaluated the hemodynamic impact of a novel STJ reconstruction surgery in a patient with a dilated ascending aorta but a normally functioning aortic valve. The surgical procedure aims to restore the sinus profiles of the aortic root and the STJ, hypothesizing that a normal-like root morphology would yield normal blood flow patterns, but this remains unquantified. While postoperative imaging could facilitate analysis of morphological differences resulting from the surgery, and hemodynamic assessment could be performed using echo and 4D-flow MRI, these can only be done in retrospect, limiting the surgeon’s preprocedural planning primarily to imaging and experience. Therefore, the current study provides cardiothoracic surgeons with a simulation-guided surgical planning framework to predict postoperative hemodynamics in the root and ascending aorta and evaluate valve function. Preplanning STJ reconstruction based on optimized model-predicted hemodynamics may mitigate the risk of complications, including root aneurysm, graft failure due to abnormal flow-induced wall stresses, and valve regurgitation, as well as the need for potential reinterventions.

We utilized patient-specific aortic geometries constructed using our previously demonstrated workflow (Section 2.2), applied physiologically relevant boundary conditions (Section 2.3), and employed a validated and widely adopted finite element solver for CFD and FSI simulations (Section 2.4). We then compared the simulated hemodynamics in the ATAA patient with those of a control subject to assess whether the new technique restores normal blood flow patterns in the root and STJ regions (Section 3). Evaluating key qualitative and quantitative hemodynamic characteristics in the ascending aorta (Section 2.5), such as the presence of coherent vortical structures in the sinuses of Valsalva, volume average velocity, kinetic energy density, viscous dissipation rate, TAWSS, and OSI, we demonstrated that the new surgical technique restores hemodynamic patterns more similar to those of the control subject in the root and ascending aortic regions, although future studies should also consider focusing on regions beyond the graft such as the aortic arch, as once the ascending region is grafted it is no longer at risk of further dilation. We also demonstrated that the FSI results largely corroborate the initial findings from the CFD analysis, although we observed some reasonable differences when incorporating valve motion.

### Morphological Characteristics of the Aortic Root

Our work builds upon previous studies of ATAA that often truncate the aortic geometry at the STJ, thereby excluding the aortic root, or that incorporate an idealized valve model rather than the one derived from patient images. Here, we developed a novel workflow to segment the aortic valve from patient images, fusing it into the aorta to perform FSI simulations (Section 2.2.1). A morphological analysis confirmed a dilated ascending aorta with a normal aortic root in the ATAA patient (Fig. 2a). The dimensions of both the aorta and the valve in the control subject were found to be much lower than those of the ATAA patient, even after intervention, consistent with the patient’s overall build. While the root dimensions were within the normal range for all three cases (Fig. 2b), the measurements for the ATAA patient are on the higher end of the spectrum and exceed the standard deviation for the preoperative non-coronary sinus-to-commissure length (L_SN_), compared to previously reported measurements [79]. At the same time, the control subject showed the greatest variance in the size of the aortic root and leaflet area, compared to the ATAA patient (Fig. 2b,c).

As the current Hegar dilator-based surgical technique is root-sparing, we expect the leaflet areas to remain the same before and after surgery. However, the differences in mean leaflet area between preop and postop cases (nearly 10%) could be partly attributed to errors aggregated from manual segmentation and model smoothing processes, although this falls within the inter-observer variability for distance and area measurements from medical images [83, 84]. At the same time, we also noted substantial differences in the LVOT shape of the ATAA patient between preoperative and postoperative imaging that was obtained approximately six months after the procedure (Fig. 8 and Appendix A). Centerline analysis suggested the Dacron graft, used to replace the removed portion of the ascending aorta, is shorter than the original vessel length (Fig. 2a). As a result, the LVOT is angled further upwards, as evidenced in the postoperative CTA images (Appendix A). We believe that this change in orientation could contribute to the altered shape of the postoperative LVOT cap. It is also unclear whether these differences in the LVOT morphology or valve geometry are due to tissue remodeling in response to the altered hemodynamic environment. The effects of these interleaflet morphological differences and the LVOT profile on the individual leaflet mobility and the ascending aortic hemodynamics should be investigated in future studies across a larger patient cohort.

### Aortic Root Hemodynamics from CFD

Examining the CFD cases on their own, we qualitatively demonstrated that the flow patterns in the region of interest for the ATAA match more closely between the postoperative and control cases, compared to the preoperative case, due to the lack of coherent vortices in the aortic sinuses in the preoperative case (Fig. 4a). The rounded sinus profile is a marker of a healthy aortic root that guides the flow through the ascending aorta; therefore, its reconstruction and narrowing of the artery at the STJ are crucial to generating the sinus vortices [85–87]. Our results are corroborated by previous studies investigating Marfan syndrome or aortic valve leaflet calcifications, underscoring the importance of these recirculating flow structures for proper valve function [88, 89].

Quantitatively, the control and postoperative cases exhibited higher average velocities (Fig. 4b) and kinetic energy (Fig. 4c), consistent with previous studies [90]. This is likely due to the reduced diameter of the waist in the STJ region, which allows for the vortices entrapped within the sinuses to pull upon the entering fluid in a centrally positioned jet, as opposed to the preoperative case, in which the inflow impinges lower along the ascending aorta, with a large flow separation zone on the inner curvature of the vessel. The narrowing of the STJ in the control and postoperative cases, and its interaction with the inflow jet, likely amplifies gradients across the shear layers, thereby increasing the rate of energy dissipation. Further, a steep increase in the viscous dissipation rate when the inflow impinges directly on the wall of the ascending aorta, which is unique to the CFD study with no aortic valve, is likely a result of the impinging of the inflow jet on the ascending aorta wall being the first contact between the fluid and any boundary.

Our results also demonstrated that the region of jet impingement on the outer curvature of the ascending aorta is also the region with the highest TAWSS (Fig. 5a). However, the magnitude of this high shear region appears to be influenced by the interaction of the jet with the STJ and the vortex on the inner curvature. Higher inflow jet velocities due to a narrowed STJ, followed by a sweeping motion of the jet constrained against the outer wall by the inner vortex, result in a thin layer with elevated velocities and shear forces on the ascending aorta. The control and postoperative cases have regions of high shear nearer the brachiocephalic artery, whereas for the preoperative case, this area of high shear is lower along the ascending aorta. For the most part, the region of high stress is confined to the outer curvature of the ascending aorta, but we observe slight discrepancies for the postoperative case, with the anterior portion showing higher shear patterns around a pocket of low shear. This shift in shear profile is likely due to the unique shape of the inlet cap or the differing LVOT angle in the postoperative case, as discussed earlier (Appendix A). When comparing the three CFD cases, we see that the postoperative and control cases exhibit higher OSI in the root than the preoperative case, suggesting greater flow reversal and recirculation in the sinuses, indicative of prominent vortices.

### Motivation for FSI and Comparison against CFD

While CFD provides detailed hemodynamic characteristics of the ascending aorta at a substantially low computational cost, and has been adopted in several studies on patient-specific modeling of ascending aorta hemodynamics [29, 32, 90], the aortic valve plays a dominant role, particularly in the root hemodynamics. Therefore, it is essential to incorporate the patient’s aortic valve into the model to elucidate its interactions with blood flow in its vicinity in the aortic root. We developed a workflow to segment the aortic valve and created a body-fitted mesh around it to perform FSI using the ALE formulation of Navier-Stokes equations. Alternative modeling approaches include creating an unfitted mesh so that the valve is simply ‘immersed’ in the background fluid domain [91–93]. The advantages and disadvantages of these approaches have been reviewed at length in other studies [94, 95]; here, we highlight only the most relevant aspects.

Despite the complexities of the manual segmentation workflow for creating a fitted unstructured mesh around the aortic valve, when coupled with ALE-based FSI, it resolves velocity gradients and pressure and shear forces with greater accuracy than unfitted approaches. This is particularly relevant during valve opening when the velocity gradients and shear are elevated. While immersed methods overcome the challenges of creating fitted meshes, these approaches typically fit a box around the immersed object [96–98], which can underresolve near-wall flow gradients; although, curvilinear immersed methods were developed to alleviate these issues [99]. Both ALE-based FSI and a class of immersed methods with unfitted meshes (e.g., ghost-cell or ghost-fluid approaches [100]) require a contact model to simulate valve closure [101, 102]. The classical continuous force-based immersed methods do not require a contact model, but they smear the valve’s influence on the blood across a diffuse interface, potentially increasing leaflet separation even in a fully closed state and underresolving near-wall gradients [103, 104]. When the valve is closed, the flow loses its momentum and kinetic energy, creating small-scale residual vortices that are typically flushed during the following opening cycle. While the shear forces are usually low during closure, the valve tissue is subjected to high stresses and strains. Therefore, in this study, we adopted the fitted-mesh approach with ALE formulation for FSI to simulate aortic valve opening.

Our FSI simulation results qualitatively corroborate the CFD findings, showing coherent vortices in the aortic sinuses for the control and postoperative cases, but not for the preoperative case (Fig. 6a). Inclusion of the leaflets dynamically changes the inflow cross-sectional area, unlike CFD where the inflow freely enters the root region, thereby increasing jet velocities, kinetic energy, and the rate of energy dissipation due to viscosity (Fig. 6b-d). Further, the gradual opening of these compliant leaflets might provide a cushioning effect, thereby dampening the steepness with which the energy dissipates, contrary to the CFD case. The shear patterns are also mostly consistent between CFD and FSI, with the high-shear region aligned with the inflow jet impingement (Fig. 7). However, the regulation and positioning of the inflow jet by the leaflets result in slightly elevated shear on the anterior wall, unlike in the CFD case. The aortic valve, therefore, acts as a flow modulator by both energizing it and streamlining flow perturbations resulting from changes in the LVOT morphology in response to intervention (Appendix A).

### Leaflet Mobility Analysis

The leaflet-opening dynamics from the FSI simulations suggest a similar GOA before and after surgery for the ATAA patient (Fig. 8). This result is not unexpected, as the root diameter and dimensions are not substantially altered post-intervention. We also note that the orifice shape is somewhat triangular, also evidenced in previous studies on image-based modeling of native valves [35, 105, 106], and is unlike the circular orifice found in prosthetic valve models [107, 108]. Additionally, the magnitudes of simulated leaflet displacements were found to be consistent with normal leaflet displacements reported in literature [35, 109]. However, the leaflet excursion analysis of the ATAA patient suggests that changes in the LVOT cross-section could affect leaflet opening, thereby altering downstream hemodynamics, which may be optimized via simulations before direct intervention.

### Effects of Tissue Heterogeneity

Our FSI model accounts for vessel wall deformation and its interaction with blood flow. In the baseline simulations presented so far, our analysis suggested that the aortic wall undergoes small deformations (max. displacement 0.6–1 mm) and strains (peak strain magnitude 0.15–0.25) when external tissue support is modeled via Robin-type boundary conditions. We have further substantiated the dominant effect of external tissue support by analyzing the model sensitivity to tissue heterogeneity.

In our baseline simulations, we assumed that the properties are homogeneous across the tissue subdomain, implying that they are uniformly distributed across the vessel wall and valve in all three cases, with a sudden transition between the valve and tissue subdomains. However, the material properties of the surgically implanted graft differ substantially from those of native tissue. Likewise, a sudden transition in the material parameters between valve and tissue is likely unphysical. To address these issues, we performed a sensitivity analysis by (i) allowing the parameters to vary between the tissue and graft subdomains (Fig. 1e) and (ii) allowing a smooth transition of the material parameters between the valve belly region and the vessel wall using a Laplace-Dirichlet solution (Fig. 1f).

We modeled graft behavior using the neo-Hookean constitutive model, similar to the native tissue (Eq. (6)), but with parameters fit to match reported values for Dacron polymer [110], resulting in an elastic modulus approximately 7.8 times larger than that of native tissue. As a result, employing graft material properties in the reconstructed ascending aorta made the vessel marginally stiffer with slightly lesser displacements and strains (Fig. B2a,b in Appendix B). The differences in mean displacements and strains with respect to the baseline case with uniform material properties were mm and 3 %, respectively. Likewise, allowing a smooth transition of the material parameters between the valve and the main arterial wall resulted in at most 0.1 mm difference in leaflet displacements and about 4 % difference in strains (Fig. B2c,d in Appendix B). These slight differences in the vessel or valvular deformation due to tissue heterogeneity did not substantially alter flow patterns in the aortic root or the ascending aorta.

### Study Limitations and Concluding Remarks

The study has several limitations that should be acknowledged. First, the coronary arteries were not included despite their proximity to the aortic root and STJ and their potential impact on the sinus vortices during valve closure [111]. However, our primary analysis of root hemodynamics was performed during systole, during which coronary drainage is low [112]. Nevertheless, we plan to enhance our FSI modeling framework to account for contact between the leaflets and to simulate leaflet dynamics and blood flow in the root throughout the cardiac cycle.

Second, some assumptions were made regarding the boundary conditions. A fixed inflow profile was applied at the inlet for both preop and postop conditions, though this is routinely performed [27, 29, 32, 33]; a multiscale coupling with a closed-loop, lumped parameter network could be used to simulate a variety of conditions, including variable heart rate and exercise [113]. Likewise, the outlet boundary condition parameters were tuned to match only postoperative data. Therefore, the analysis is not strictly ‘patient-specific’ and focuses on analyzing the effect of geometrical variations due to surgical intervention; however, we made this assumption to minimize the influence of boundary conditions and other model parameters on the hemodynamic assessment. Further, BSA scaling was applied to these inflow and outflow boundary conditions to enable a meaningful comparison against the control subject. Neither CFD nor FSI models account for the interaction between the left ventricle and the root. The root could undergo substantial translational motion as the left ventricular myocardium contracts and exerts a pulling force on the root via the LVOT [114]. We will evaluate the effect of these assumptions on the boundary conditions in a follow-up investigation.

Third, FSI simulations were performed only during systole and not the whole cardiac cycle. Therefore, our analysis was limited to the first cycle and, unlike CFD, was not continued over multiple cycles to achieve limit-cycle convergence. In an ongoing project, we are developing robust methods to simulate contact between the leaflets and will then extend the analysis throughout the cardiac cycle [115, 116].

Fourth, we employed the isotropic, hyperelastic neo-Hookean material model for the tissue domain. Biomechanical experiments have shown that the aorta and valve exhibit anisotropic, hyperelastic material responses [117]. However, when the vessel is constrained by external tissue support, the tissue strains were low (0.15–0.25), to justify linear and isotropic behavior. We are developing rule-based methods to generate circumferential and longitudinal fiber directions for patient-derived valve models [118, 119]. Together with the contact model, we will evaluate the effects of these constitutive models on leaflet opening and closing dynamics in a follow-up study.

Fifth, in this study, we had access only to single-phase, retrospective CTA images obtained during diastole. As a result, we could not validate the predicted shape of the leaflet orifice against image data. While our simulated open valve state agrees with prior *in vivo* aortic valve analysis [120], we will validate our FSI modeling workflow in a cohort of prospectively selected patients who have undergone postoperative dynamic CTA and will use 4D-flow MRI data for hemodynamic validation.

Lastly, although the gated CTA was acquired *in vivo* during diastole, we assumed that the initial segmented model represents the reference, stress-free configuration within the adopted nonlinear mechanics framework for FSI. Although we have prestressed the aorta in our prior work on aortic dissection [21], it is not straightforward to prestress the vessel wall with a fully integrated valve, and it is part of our ongoing work. Moreover, given the small overall deformation with external tissue support, we believe prestressing will not have a major impact on the blood flow patterns, although it could alter the magnitude of tissue stresses and strains.

In conclusion, we have evaluated the hemodynamic impact of a Hegar dilator-based, root-sparing, ascending aortic aneurysm reconstruction surgery on the local hemodynamics. In a retrospectively selected patient with preoperative and postoperative CTA imaging, models of the thoracic aorta and aortic valve were constructed using a novel workflow. CFD and FSI simulations were performed using a previously validated, widely adopted multiphysics finite element solver, with physiologically relevant boundary conditions applied. The workflow is repeated on a control subject with BSA-matched boundary conditions. Simulated hemodynamics were compared across all three cases (preoperative, postoperative, and control) to assess whether the procedure’s ability to restore normal root morphology would also restore normal hemodynamics. Notably, both CFD and FSI simulations exhibited similar hemodynamic characteristics between the postoperative and control cases, although the FSI data showed greater magnitude differences than CFD. Simulated hemodynamics were also found to be less sensitive to the graft material parameters and leaflet material heterogeneity. While the majority of these patients currently undergo size-based clinical evaluation, our analysis shows that, when addressing cases of STJ dilation, the shape of the reconstruction is equally important, including the STJ’s constrictive waist and the angle of the LVOT, rather than just its size. Although the analysis is retrospective, the modeling framework could be applied pre-surgery and may assist in graft design to yield optimal ascending aortic hemodynamics and valve function; however, further validation is needed in a larger patient cohort.

## Abbreviations

ATAA: Ascending Thoracic Aortic Aneurysm
BSA: Body Surface Area
CFD: Computational Fluid Dynamics
CTA: Computed Tomography Angiography
FEA: Finite Element Analysis
FSI: Fluid-Structure Interaction
GOA: Geometric Orifice Area
LCC: Left Coronary Cusp
LVOT: Left Ventricular Outflow Tract
NCC: Non-Coronary Cusp
OSI: Oscillatory Shear Index
RCC: Right Coronary Cusp
STJ: Sino-Tubular Junction
TAWSS: Time Average Wall Shear Stress

## Acknowledgments

HZ would like to acknowledge a graduate research fellowship through the US National Science Foundation. VV and HT would like to acknowledge partial funding support from Columbia University SEAS Interdisciplinary Research Seed (SIRS) grant / Blavatnik Fund. HT would like to acknowledge funding from the Rudin Foundation. We also acknowledge computing resources from ACCESS (SDSC Expanse) and the Columbia Ginsburg HPC cluster for supporting this work.

## Declarations

### Human Subjects

This single-center study was performed following HIPAA-compliant processes and guidelines and was approved by the Columbia University Medical Center Institutional Review Board (IRB#AAAR2949).

### Consent for Publication

All authors have agreed with the content and given explicit consent to submit the manuscript for publication.

### Data Availability

Data will be shared upon a reasonable request from the corresponding authors.

## Appendix A

**LVOT orientation pre and post-surgery**

**Fig. A1.**
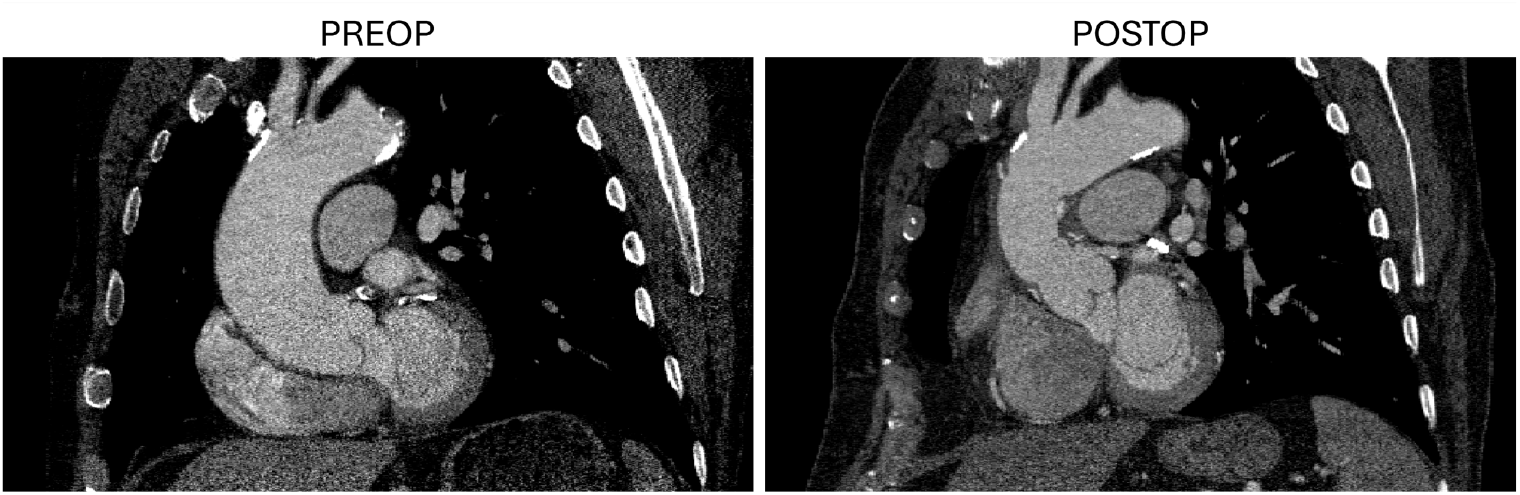
A coronal view of (left) preoperative and (right) postoperative CTA images in the ATAA patient showing differences in LVOT orientation after intervention.

A direct comparison of the ATAA patient’s preoperative and postoperative CTA images in a coronal view shows marked differences in LVOT orientation (Fig. A1). In the preoperative case, we see an inferiorly leaning LVOT compared to the obliquely oriented LVOT post-surgery. This change in angle is likely due to the graft in the ascending region being shorter than the dilated vessel it replaced. As a result, the root is pulled upward, leading to a more direct flow from the ventricle. Further, the postoperative imaging was obtained approximately 6 months after the procedure, suggesting that changes in LVOT and aortic root morphology may be due to tissue remodeling in response to altered hemodynamic loading and surrounding structures.

## Appendix B

**Effect of Tissue Heterogeneity**

We considered two cases of tissue heterogeneity and evaluated their effects on ascending aortic hemodynamics, comparing them with the baseline case presented in Section 2.4. Notably, the baseline case employed uniform homogeneous material properties throughout the aortic tissue and aortic valve. Therefore, in one instance, we employed Dacron material properties in the vessel region that was replaced with the graft. In another case, we created a transitional layer that smoothly transitions the material properties from the valve leaflets to the main arterial wall.

Applying Dacron material properties resulted in slight differences in displacement, with a local time-averaged maximum of 0.3 mm and a spatially-averaged maximum of 1 mm (Fig. B2a). Differences in the corresponding strains were within 3–4 % (Fig. B2b). Likewise, varying the material parameters smoothly between the leaflet belly and the main arterial wall resulted in negligible differences in local time-averaged leaflet displacements (∼0.1 mm) and spatially-averaged maximum displacement (∼1.4 mm), as shown in Fig. B2c. Differences in the corresponding strains were similarly kept low under 4–6 % (Fig. B2d). Importantly, the slight differences in the tissue deformation due to these material property variations did not substantially alter blood flow patterns in the aortic root or the ascending aorta.

**Fig. B2.**
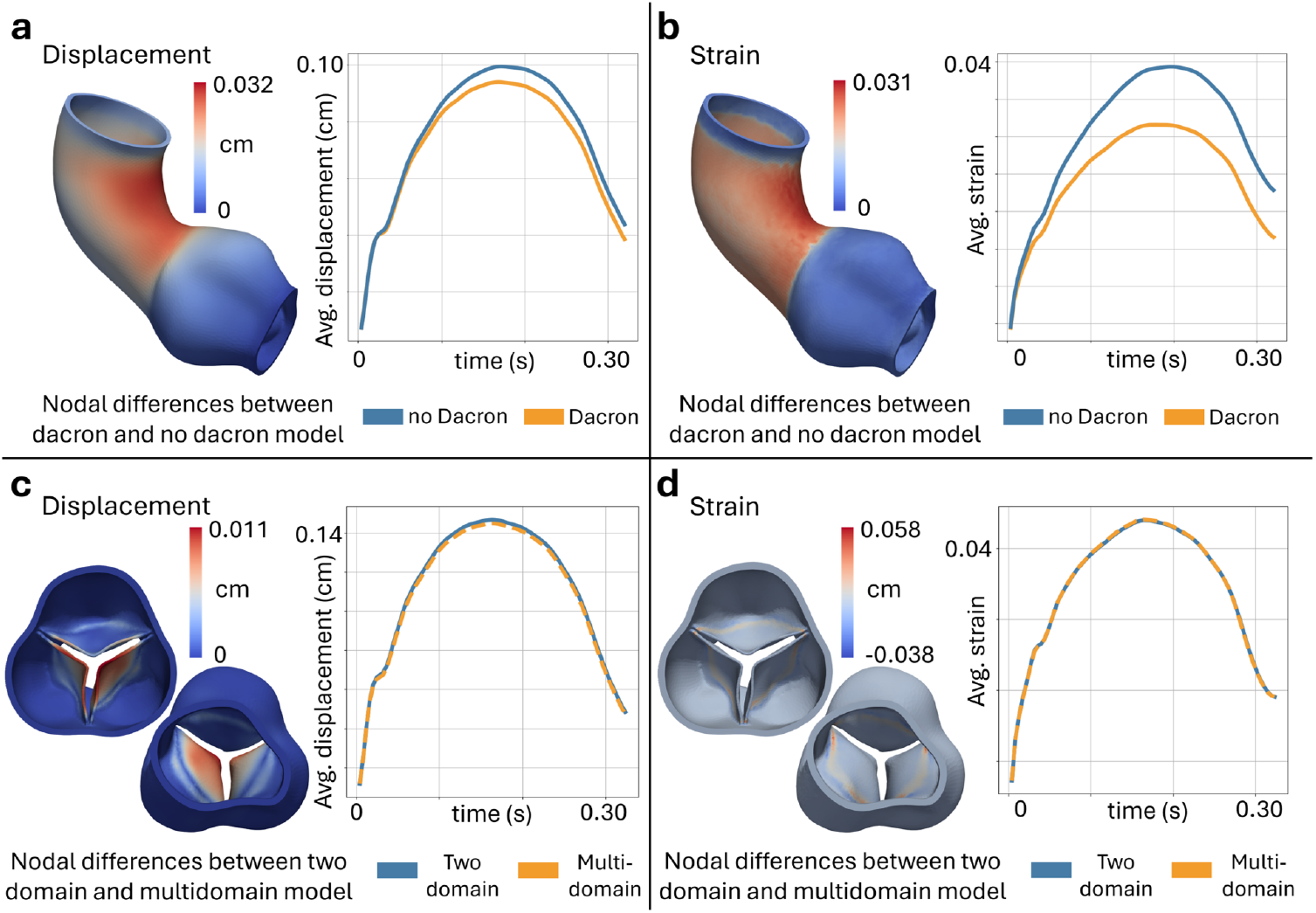
Comparison of (left) nodal displacements and (right) strain magnitude between the baseline model and (top) employing Dacron-based graft material properties in place of the native vessel in the reconstructed ascending aortic region, and (bottom) allowing a smooth transition of the material properties from the leaflet belly to the main arterial wall using a multi-domain model. The surface map displays time-averaged differences in nodal displacements or strains between the sensitivity case and the baseline model. The line plot displays spatially-averaged differences in nodal displacements or strains, plotted as a function of time, during systole.

## Footnotes

1 The aortic root is located at the base of the aorta, housing the aortic valve, the sinuses, and the coronary ostia.

2 https://github.com/SimVascular/svFSI

